# Tissue-engineered models of lung cancer premalignancy

**DOI:** 10.1101/2023.03.15.532835

**Authors:** Rachel Blomberg, Kayla Sompel, Caroline Hauer, Brisa Peña, Jennifer Driscoll, Patrick S. Hume, Daniel T. Merrick, Meredith A. Tennis, Chelsea M. Magin

## Abstract

Lung cancer is the leading global cause of cancer-related deaths. Although smoking cessation is the best preventive action, nearly 50% of all lung cancer diagnoses occur in people who have already quit smoking. Research into treatment options for these high-risk patients has been constrained to rodent models of chemical carcinogenesis, which are time-consuming, expensive, and require large numbers of animals. Here we show that embedding precision-cut lung slices within an engineered hydrogel and exposing this tissue to a carcinogen from cigarette smoke creates an *in vitro* model of lung cancer premalignancy. Hydrogel formulations were selected to promote early lung cancer cellular phenotypes and extend PCLS viability up to six weeks. In this study, hydrogel-embedded lung slices were exposed to the cigarette smoke derived carcinogen vinyl carbamate, which induces adenocarcinoma in mice. At six weeks, analysis of proliferation, gene expression, histology, tissue stiffness, and cellular content revealed that vinyl carbamate induced the formation of premalignant lesions with a mixed adenoma/squamous phenotype. Two putative chemoprevention agents were able to freely diffuse through the hydrogel and induce tissue-level changes. The design parameters selected using murine tissue were validated with hydrogel-embedded human PCLS and results showed increased proliferation and premalignant lesion gene expression patterns. This tissue-engineered model of human lung cancer premalignancy is the starting point for more sophisticated *ex vivo* models and a foundation for the study of carcinogenesis and chemoprevention strategies.

## Introduction

Lung cancer is a serious global health problem, and for years has been the leading cause of cancer-related deaths across the world^1^. Despite advances in targeted agents and immunotherapies, there remains a need to develop new prevention options for lung cancer patients^2^. The clearest form of lung cancer prevention is to reduce smoking prevalence, but more than half of all lung cancer cases are diagnosed in former smokers. In fact, a new deep learning model can now accurately predict lung cancer risk in former smokers from a single computed tomography (CT) scan^3^. This innovation underscores the need for interventions targeting premalignant lesions in these high-risk individuals. Despite the tremendous effort to translate lung cancer therapies to the clinic, 5-year survival for lung cancer is still only 22%.^4^ Chemoprevention holds the potential to prevent tumor formation and provides options for patients to minimize lung cancer risk. Clinical translation of chemoprevention could be accelerated by preclinical models that enable studies of lung premalignant lesion biology and prevention mechanisms. *In vivo* mouse models are currently the gold standard for preclinical research in lung cancer^5-7^, since observing cancer in an intact organism allows researchers to fully recapitulate the kind of complex multicellular interactions that underlie cancer initiation and progression. Combinatorial genetically-engineered mouse models, such as p53 loss in conjunction with K-Ras mutation, can produce rapidly developing tumors that recapitulate a range of human lung cancer stages^8^. Likewise, susceptible strains of mice can be exposed to cigarette smoke or smoke components like urethane (ethyl carbamate) or vinyl carbamate to model chemical carcinogenesis of lung cancer caused by smoking. These models directly relate to the carcinogenesis process in humans and recapitulate human lesions across the spectrum of progression^9^. A key advantage of chemical models compared to genetically engineered models is the opportunity to test the prevention approaches required to ensure high-risk individuals, such as former smokers, have ways to minimize cancer risk. New *et. al.*, for example, demonstrated that murine adenomas express markers of epithelial to mesenchymal transition (EMT) and that this shift toward EMT could be reversed by increased prostacyclin^10^ using urethane exposure in mice. Similarly, Tennis *et. al.* showed the potential of the prostacyclin analog iloprost, a PPARγ agonist, as a chemoprevention agent in the urethane mouse model of early lung cancer^11^. Although *in vivo* models have provided insights into the mechanisms underlying cancer initiation, progression, and treatment, such models tend to be expensive and low-throughput. New tools and technologies that enable us to develop effective chemoprevention strategies are critical to reducing mortality from lung cancer.

Precision-cut lung slices (PCLS) are a well-established technology with great potential for *ex vivo* modeling of the complex tissue interactions in lung cancer^12^. PCLS are generated by filling lung tissue with agarose to provide stability and then sectioning on a vibratome to generate intact tissue slices which can be maintained in standard tissue culture conditions^13,14^. One major advantage of PCLS over more reductionist systems, including organoid culture, is the presence of multiple cell types and lineages preserved in the native architecture^12,15^. Lung morphology, alveolar space, arterioles and veins, enzymatic activity, and cellular functions are kept intact in these thin slices of lung tissue cultured *ex vivo*.^16^ PCLS have enabled studies of acute lung conditions such as response to bacterial or viral infections^12^, but the utility of PCLS to model chronic lung conditions has been limited by short viability in culture (7 to 10 days)^17,18^. Therefore, use of PCLS to investigate lung cancer has been limited. Two studies generated PCLS from human and murine lung tissue with tumors to test therapeutic approaches over one or seven days.^19,20^ A recent study generated PCLS from *in vivo* chemical and genetic models of carcinogenesis and showed recapitulation of premalignancy and chemoprevention signaling over 7-10 days.^21^ Such studies suggest PCLS are a promising technology for high throughput chemoprevention drug screening and efficient *in vivo* studies^22-27^ if *ex vivo* viability and functionality could be extended.

Hydrogel biomaterials are powerful tools for studying the interactions between cells and the tumor microenvironment^28,29^. Both natural and synthetic biomaterials can be used to encapsulate cells in an environment that mimics critical biochemical and/or biomechanical cues from the native tissue environment. Two commonly used natural biomaterials are collagen I, a primary component of fibrillar ECM, and Matrigel, a murine tumor-derived ECM extract rich in basement membrane components like laminin. Culturing murine tumor fragments in naturally derived materials, Matrigel and collagen, showed that collagen I enhanced tumor-progressive phenotypes, specifically integrin-mediated cellular invasion^30^. These data agree with observations that collagen I expression increases during cancer progression^31,32^ and that collagen architecture can strongly influence cancer cell behavior^33,34^, demonstrating the importance of natural ECM components to tumor progression. Synthetic hydrogels provide a more precisely tunable platform than natural hydrogels, particularly in terms of mechanical properties. Gill et. al. embedded a metastatic lung cancer cell line in a PEG-based hydrogel and by modulating the elastic modulus and adhesion peptide (RGD) concentration observed changes in the organization and polarity of self-forming cellular structures in 3D culture^35^. The results in this 3D model align with studies of human tumors that showed tumor progression correlated with increased tissue stiffness^36^. Lewis et al. demonstrated in 3D co-culture within PEG-based hydrogels that a cancerous lung epithelial cell line (A549) induced faster fibroblast migration rates and increased matrix metalloproteinase (MMP) activity^37^. Rigorous research in biomaterials-based models has established the value of modeling multicellular interactions within a physiologically relevant 3D environment for studying lung cancer progression and metastasis^38^, however, less has been reported about modeling lung cancer premalignancy *ex vivo*.

Our group pioneered a technique for prolonging PCLS viability and maintaining normal tissue architecture for at least three weeks in culture by embedding the tissue within a hydrogel biomaterial^39^. These hydrogels were comprised of a polyethylene glycol (PEG) norbornene backbone and crosslinked with dithiothreitol via cytocompatible click reaction. Cellular adhesion peptides mimicking fibronectin (CGRGDS) and laminin (CGYIGSR) were incorporated to allow the embedded tissue to attach to and interact with the hydrogel. PCLS were embedded using a dual-layer approach in custom molds to create 3D constructs with lung tissue surrounded by hydrogel on all sides and cultured without significant decreases in viability or changes in structure for three weeks^39^. In this study, we present a tissue-engineered model of lung cancer premalignancy in hydrogel-embedded PCLS. Assessments of metabolic activity and cellular viability indicated that hydrogel embedding maintained cells within murine-derived PCLS for an unprecedented six weeks in culture. A design of experiments approach selected a biomaterial microenvironment that best supported induction of chemical carcinogenesis *ex vivo.* Quantification of proliferation, tissue morphology, and gene expression demonstrated that treatment with the chemical carcinogen vinyl carbamate induced pre-malignant phenotypes. Additionally, chemoprevention agents were diffused through the hydrogel and induced gene expression changes known to be protective against tumorigenesis in hydrogel-embedded PCLS. Finally, the design parameters established using murine PCLS were validated using human lung tissue. Collectively, these results present hydrogel-embedded PCLS as a new model for lung carcinogenesis.

## Methods

### Animal Use

All animal use was carried out in accordance with protocols approved by the Rocky Mountain Regional VA Medical Center Veterinary Medical Unit (VMU) Institutional Animal Care and Use Committee. Wild-type female A/J mice^21,40,41^ were purchased from Jackson Laboratories at eight weeks old and acclimated to standard housing at the VMU for one to two weeks before harvest for PCLS. For atomic force microscopy, lungs were harvested from mice exposed to urethane or saline. At 10 weeks of age, female mice were injected IP with 1mg/kg urethane or saline and weighed daily for the first week and weekly thereafter to monitor toxicity. Mice were then harvested for PCLS at 16 weeks after urethane injection.^40,42^

### Generation of PCLS

Murine: 8–10-week-old mice with no exposures to chemical carcinogens or 16-week urethane/saline-exposed mice were sacrificed by a lethal dose of Fatal Plus (Vortech Pharmaceuticals, Ltd.), delivered by intraperitoneal injection, and secondly by exsanguination. The lungs were cleared with a cardiac perfusion of 10 ml of sterile phosphate-buffered saline (PBS; Gibco) through the right ventricle. Immediately after, the lungs were filled via tracheal perfusion with 1 ml of 1.5% low melting point agarose (Fisher Scientific) dissolved in sterile HEPES buffer. The whole mouse was placed on ice for 10 min before the lungs were extracted and placed in 1 ml of DMEM media with 0.1% Penicillin/Streptomycin/Fungizone (P/S/F) antibiotics/antimycotics. The lobes were then sliced into 500 mM whole slices with a vibratome (Campden Instruments) at 12 mm/s speed. A biopsy puncher (4 mm, Fisher Scientific) was used to create standardized punches. The punches were then washed with media 3 times, with 30-minute incubations at 37°C between each wash to remove the agarose. Punches were cultured overnight in 24 well plates before embedding.

Human: Agarose-filled lung cores^43^ (Deidentified by provider, 20yr old male, non-smoker, Lung # 495, UNOS: AJHF491) were generously provided by the laboratory of Dr. Hong Wei Chu (National Jewish Health) and subsequently sliced, punched, and cultured as described above.

### PEG-Norbornene (PEG-NB) synthesis

Eight-arm 10 kg/mol PEG-hydroxyl (PEG-OH; JenKem Technology) was conjugated with norbornene as described previously^44,45^. Lyophilized PEG-OH (5 g) and 4-Dimethylaminopyridine (DMAP, 0.002 mol; Sigma-Aldrich) were dissolved in anhydrous dichloromethane (DCM; Sigma-Aldrich) and then Pyridine (0.02 mol; Fisher Scientific) was injected to the mixture. In a separate reaction container, N,N’-Dicyclohexylcarbodiimide (DCC, 0.02 mol; Fisher Scientific) was dissolved in DCM with Norbornene-2-carboxylic acid (0.04 mol; Sigma-Aldrich). This mixture was stirred at room temperature for 30 minutes, filtered through Celite 545 (EMD Millipore), and the filtrate then added to the PEG-OH mixture and allowed to react for 48 hours while protected from light. The product was then sequentially washed by phase separation in 5% sodium bicarbonate, brine, and distilled water. The organic phase was concentrated on a rotary evaporator and the product precipitated with diethyl ether (Sigma-Aldrich) at 4°C overnight. The product was then purified by dialysis in 1 kg/mol MWCO tubing (Repligen) against deionized water for three days, with water changes every 24 hours. The final resulting product was lyophilized and then characterized by 1H NMR (Bruker DPX-400 FT NMR spectrometer, 300 MHz) to ensure at least 90% functionalization.

### Rheological characterization of hydrogels

The elastic modulus of hydrogels was assessed using parallel plate rheology according to established methods^46,47^. Hydrogels were cast into discs with 1-mm height and 8-mm diameter and equilibrated in PBS overnight. The samples were then loaded onto an 8-mm diameter parallel plate geometry on a rheometer (Discovery HR2, TA instruments). The geometry was lowered to apply 0.03 N axial force and this gap distance recorded. Measurements were taken at increasing levels of compression (20%, 30%, and 40%) identify the compression level where the storage modulus (G’) plateaued. Values taken at 30% compression are reported here. Samples were then subjected to a range of frequency oscillatory strains from 0.1 to 100 rad/s at 1% strain. The elastic modulus (E) was calculated using rubber elasticity theory, assuming a Poisson’s ratio of 0.5 for bulk measurements of the elastic hydrogel polymer network.

### Hydrogel embedding and culture of PCLS

The 4-mm PCLS punches were embedded in a silicone mold with 8-mm diameter wells as described previously^39^. First, a bottom layer of hydrogel solution was polymerized using a light-mediated thiol-ene reaction. Hydrogel solutions consisted of PEG-NB macromer (5 or 7 wt%), DTT crosslinker (r ratio = 0.9; Fisher), and the cellular adhesion peptides CGRGDS (0.1 mM), CYIGSR (0.2 mM), and CGFOGER (0.1 or 1 mM; GL Biochem) in culture media (Figure 1B). The photoinitiator lithium phenyl-2,4,6-trimethylbenzoylphosphinate (LAP; Sigma Aldrich) was added for a final concentration of 2.2 mM. Next 25 µl per well of the hydrogel precursor solution was pipetted to fill the well. The solution was exposed to 365 nm UV light at 10 mW/cm^2^ for 5 min to facilitate crosslinking. Then a single PCLS punch was laid flat and centered on this base layer in each well, covered with an additional 25 µl of hydrogel solution, and polymerized under UV light using the same conditions as listed above. Hydrogel formulations that were intentionally designed to be slightly off-stoichiometry (thiol:ene = 0.9) enabled covalent attachment of the second layer to the first during the second polymerization step. Flexible plastic scoops were used to gently lift and transfer each hydrogel-embedded PCLS punch into the well of a 24-well plate.

**Figure 1:**
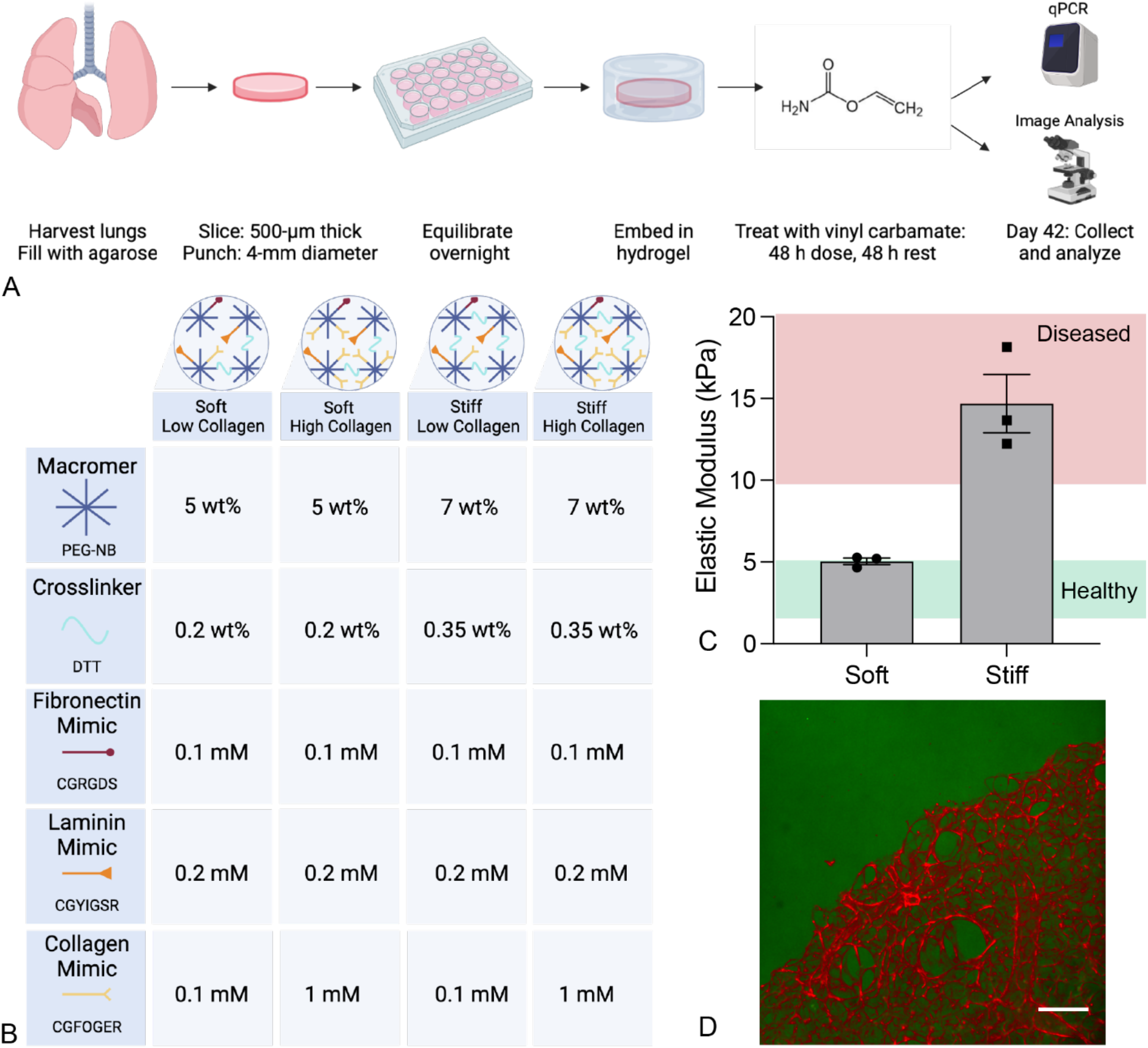
PEG-NB hydrogels were tuned to recapitulate key features of the premalignant microenvironment. A) Schematic of the experimental design for generating tissue-engineered models of lung cancer premalignancy in hydrogel-embedded PCLS. B) The compositions of hydrogel formulations tested to support *ex vivo* premalignancy development. C) Rheology results for soft and stiff hydrogel formulations revealed that these materials mimic the mechanical properties of healthy (1-5 kPa) and diseased (>10 kPa) human lung tissue, respectively. Soft hydrogels had an elastic modulus of 5.05 ± 0.20 kPa while stiff have a modulus of 14.69 ± 1.78 kPa (N=3). D) Maximum intensity projection of a confocal image. Hydrogel-embedded murine PCLS showing lung tissue labeled with CellTracker (red) and hydrogel labeled with a maleimide-conjugated AlexaFluor488 (green). Scale bar, 200 μm.

Hydrogel-embedded PCLS punches were maintained at 37°C with 5.0% CO_2_ in DMEM:F12 (1:1; Gibco) media with 0.1% fetal bovine serum (Gibco) and 0.1% 100X P/S/F antibiotics. Media was changed every 48 hours. In carcinogen experiments, cells were exposed to either 10 µg/ml or 50 µg/ml vinyl carbamate (Toronto Research Chemicals), or the vehicle control (1X sterile PBS). In chemoprevention experiments, PCLS cultures were treated with 10 µM Iloprost (Cayman Chemical) or 9 mM methyl acetate vehicle control (Sigma Aldrich) every 48 hours, or 8 μM curcumin (Cayman Chemical) or sterile water vehicle control every 24 hours.

### Presto blue metabolic activity assay

Presto blue reagent (Fisher Scientific) was diluted 1:10 in culture media and 500 µl was added to each well containing one hydrogel-embedded PCLS punch. The hydrogel-embedded PCLS punches were incubated with Presto blue solution at 37°C for 2 hours. Following incubation, triplicate samples of 150 μl of Presto blue solution were transferred to a clear 96-well plate. The fluorescence intensity of each well and empty controls was measured at 520 nm excitation on a Glomax plate reader (Promega). The average blank control values for each plate were subtracted from individual fluorescence intensity values and weekly measurements were normalized to the initial reading.

### Hydrogel-embedded PCLS imaging

To visualize both PCLS and hydrogel, tissue slices were labeled prior to embedding by incubating for 30 min at 37° in 10 µM CellTracker Orange (Thermo Fisher) in culture media. The hydrogel was labeled by incorporation of 20 µM AlexaFluor 488 C_5_ Maleimide (Thermo Fisher) during polymerization. Images of the hydrogel-embedded PCLS were acquired on a spinning-disk confocal microscope (Leica) and are presented as a maximum projection of a z-stack.

### Live/Dead staining

Live hydrogel-embedded PCLS punches were incubated with a 1:1000 dilution each of CytoDye 488 and propidium iodide (EMD Millipore, QIA76) in culture media for 40 min at 37°C. Samples were then washed once with PBS before being stored in 0.1% FBS in PBS for imaging. Image acquisition was performed on a confocal microscope (3I MARIANAS Spinning Disk) by taking 50 µm z-stacks with the 10x objective. Viability was quantified from images in Fiji (ImageJ) by generating a maximum projection for each channel, thresholding, and counting particles to determine the number of nuclei per field of view. The number of nuclei positive for propidium iodide (dead) was divided by the number stained with CytoDye 488 (total) to determine fraction of dead cells. Data are presented as the inverse: percent live cells.

### EdU incorporation assay

A 5-ethynyl-2′-deoxyuridine (EdU) incorporation assay was performed according to the manufacturer’s protocol (Invitrogen). Briefly, samples were incubated in 10 μM EdU in complete culture media for 16 hours. Samples were then fixed with 4% PFA, permeabilized with 0.5% Triton X-100, and then incubated in Click-iT reaction cocktail for 30 min. Nuclei were counterstained with 5 μM Hoechst 33342. Samples were imaged on an Olympus BX63 microscope by acquiring a z-stack over 50 µm at 10x magnification. Weiner deconvolution was performed on all z-stacks, and then image analysis performed in Fiji (ImageJ) on a maximum projection. Single channel images were thresholded to exclude background and number of nuclei determined by counting particles. The number of EdU+ nuclei was divided by the number of total (Hoechst+) nuclei to calculate percent proliferating cells.

### RT-qPCR

For each sample, three to five hydrogel-embedded PCLS punches of the same experimental group were pooled together, and RNA was extracted with the RNeasy Plus kit (Qiagen). All primers were qPCR Prime PCR Assays (Bio-Rad), and the unique IDs for each primer set are provided in Supplemental Table 1. qPCR was conducted using the standard protocol for SsoAdvanced SYBR Green Master Mix (Bio-Rad) on a CFX96 Touch (Bio-Rad). All gene expression data were normalized to the reference gene RPS18 and fold changes were calculated using the 2-ýýCt method.

### Cryosectioning

Hydrogel-embedded PCLS punches were fixed in 4% paraformaldehyde in PBS (Electron Microscopy Sciences) for 30 min, and then excess aldehyde was quenched with 100 mM glycine (Fisher) for 30 min, both at room temperature with rocking. Samples were then washed three times with PBS and excess hydrogel trimmed from around the tissue. Samples were placed in optimal cooling temperature (OCT) compound for 48 hours at room temperature to allow complete perfusion of OCT through the remaining hydrogel and to the tissue. Samples in OCT were transferred to 10×10×5 mm cryomolds (Sakura), with 3-5 samples stacked into one mold. Stacked samples were then flash frozen by submersion in liquid nitrogen-cooled 2-methylbutane (Sigma-Aldrich) and stored at -80°C until cryosectioning.

Cryosectioning was carried out on a Leica CM1850 cryostat. Frozen blocks were removed from the mold and mounted with one side pressed to the specimen disc, such that each cryosection would contain parallel cross-sections of the stacked PCLS punches. Serial sections of 10 µm thickness were acquired at –20°C and stored at -80°C for further processing.

### Atomic Force Microscopy

Mechanical properties of the tissues were analyzed as previously reported^48-51^ using a NanoWizard® 4a (JPK Instruments). Lung lobes were harvested from either healthy mice or mice bearing urethane-induced adenomas produced as previously described, fixed for 30 min in 4% PFA, and frozen in OCT, consistent with sample preparation for hydrogel-embedded PCLS. 10-µm frozen sections of healthy, adenoma, or carcinogen-treated, hydrogel-embedded PCLS were analyzed. Slides were rinsed in PBS for 15 min to remove OCT and any debris from the tissue. Tissue stiffness was determined using quantitative imaging (QI) mode with a qp-BioAC-1 (NanoandMore) cantilever with a force constant in the range of 0.15 to 0.55 N/m. Calibration of the cantilever was performed using the thermal oscillation method prior to each experiment.

Regions of interest were identified by morphology on phase-contrast images and then scanned using a set point of 5 nN and a Z-length of 2 µm. Each scan was composed of 65,536 force curves. In JPK software, the Hertz model was used to determine the mechanical properties of the tissues, and a correction for an offset in the height data was performed line by line using the JPK data processing operation. The experimental data were analyzed in GraphPad Prism software using a Brown-Forsythe ANOVA test with Dunnett’s T3 multiple comparisons, due to the significant differences in variance between the groups.

### Hematoxylin and eosin staining

Frozen slides were first brought to room temperature and then OCT cleared by dipping in PBS. Sections were fixed by submersion in ice-cold acetone for 15 min and then equilibrated for 3 min in water. Slides were then sequentially submerged in the following solutions: Hematoxylin (Richard-Allan Scientific), 1 min; DI water, dip; DI water, 1 min; 1% HCL in 95% ethanol, dip; 0.1% sodium bicarbonate, 1 min; 95% ethanol, 1 min; Eosin-Y (Richard-Allan Scientific), 1 min; 95% ethanol, dip; 95% ethanol, 1 min, 100% ethanol, 2 x 5 min; SafeClear II (Fisher) 2 x 5 min. Sections were mounted in Cytoseal 60 (Thermo Scientific) under a 24 x 50 mm coverslip before imaging. Images were blindly assessed by an expert lung pathologist for evidence of tumor lesion morphology or other abnormalities.

### Immunofluorescence staining

Frozen sections were thawed to room temperature and then equilibrated for 3 min in PBS. Sections were circled using an ImmEdge hydrophobic pen (Vector Laboratories) and blocked for 45 minutes in 5% BSA in PBS. For stains involving a mouse antibody, an additional 45-min block was performed in mouse-on-mouse blocking reagent (Invitrogen). Sections were incubated with primary antibodies (Supplementary Table S2) in 5% BSA overnight at 4°C, and then washed three times (5 minutes each wash) in 0.1% Tween-20 in PBS (PBST). Sections were incubated with secondary antibodies (5 µg/ml) in 5% BSA in PBS for 1 h at room temperature and then washed three times in PBST. For stains including lectin to visualize tissue architecture, tomato lectin (Vector Labs) was diluted 1:250 and mixed with secondary antibodies, and samples were incubated for 45 min at 37° before washing. Sections were then incubated in 300 mM DAPI for 15 min at room temperature and then washed once in PBS. Sections were mounted under a 24 x 50 mm coverslip in ProLong Gold antifade reagent (Invitrogen) and allowed to cure overnight before imaging.

### Peroxisome Proliferator Activated Receptor gamma (PPARγ) Activity Assay

After six days in culture with iloprost, PPARγ activity was assessed using a PPARγ response element (PPRE) luciferase assay. Immortalized Human Bronchial Epithelial Cells (HBEC) cells were seeded at 5000 cell/well in RPMI media with 10% FBS in a 96 well plate. The plate was incubated for 24 hours at 37°C, 5.0% CO_2_. Wells were transfected with 45 ng PPRE luciferase (a gift from Bruce Spiegelman; Addgene plasmid #1015) and 5 ng renilla control reporter vector (Promega), using TransIT-X2 transfection reagent (Mirus Bio) according to the manufacturer’s protocol. Luciferase mock and empty transfection controls were included. 15 µL of media was collected from day seven PCLS punches 24 hours after the last treatment with iloprost or vehicle control and was added to HBEC cells 24 hours after transfection with PPRE luciferase. The plate was incubated for 24 hours with PCLS media, then luciferase activity was measured using the Dual-Luciferase Reporter assay kit (Promega) on the Glomax instrument (Promega). PPRE firefly activity was normalized to renilla activity and analyzed relative to vehicle controls.

### Statistical Analysis

Experiments with two groups and a single independent variable were assessed by two-tailed student’s t-test. Dose responses with three or more groups and a single independent variable were assessed by one-way ANOVA with a Tukey’s test for multiple comparisons. Experiments with four or more groups and two independent variables were assessed by two-way ANOVA with a Tukey’s test for multiple comparisons. Data that showed significantly different variances between groups was assessed with appropriate modifications where noted. All statistical analyses were conducted in GraphPad Prism software (GraphPad Software, Inc., San Diego, CA) Hydrogel formulations were determined by a custom design of experiments generated with JMP software (SAS, Cary, NC). The resulting proliferation and gene expression data were entered back into the custom design and analyzed with a least-squares regression model to determine the most significant factors and plot the predicted response.

## Results

### PEG-NB hydrogels recapitulated key aspects of the tumor microenvironment

To engineer a PCLS-based, *ex vivo* model of lung cancer premalignancy, PCLS were generated according to standard protocols. Intact mouse lungs were filled with agarose via the trachea to provide stability during cutting on a vibratome. Thin slices (500 µm) were sectioned, and a 4-mm diameter circular biopsy punch was used to cut out uniform shapes for embedding. The punches were allowed to rest overnight in complete culture media. Embedding was performed using custom silicone molds with 8-mm diameter wells as previously described^39^. An initial base layer of hydrogel was cast and polymerized, with a ratio of macromer to crosslinker designed to leave free reactive functional groups to covalently bind the top and bottom layers of hydrogel. The PCLS punch was laid in the center of this base layer, covered with a top layer of hydrogel mixture, and polymerized. The free reactive groups on the base layer were then able to crosslink with the upper layer, providing a solid structure surrounding the PCLS punch on every side. These hydrogel-embedded PCLS punches were then cultured free-floating in standard cell culture media, with or without chemical supplementation, until the time of analysis (Fig 1A). The hydrogels were comprised of a norbornene-conjugated, eight-arm PEG macromer as the backbone, with a DTT crosslinker and a cocktail of pendant peptides designed to facilitate cellular adhesion to and interaction with the hydrogel. A design of experiments statistical approach examined key aspects of the developing tumor microenvironment, specifically increasing stiffness and collagen deposition. Varying macromer concentration (weight percent) adjusted the elastic modulus or stiffness of the microenvironment while varying concentrations of the collagen mimic peptide CGFOGER replicated increases in collagen observed in premalignant tissues (Fig 1B). Hydrogels were designed to recapitulate either healthy (1-5 kPa) or diseased (>10 kPa) lung tissue^52,53^, with soft hydrogels displaying an elastic modulus of 5.05 ± 0.20 kPa and stiff hydrogels at 14.69 ± 1.78 kPa (Fig 1C). Microscopically, hydrogel-embedded PCLS showed interactions with the hydrogel along the edge of each sample, and preserved tissue integrity throughout (Fig 1D).

### Vinyl carbamate treatment induced premalignant phenotypes in hydrogel-embedded murine PCLS

Dose testing was performed with the cigarette-smoke carcinogen vinyl carbamate, the carcinogenic metabolite of urethane, to screen for any toxicity in hydrogel-embedded PCLS during culture. Media on the hydrogel-embedded PCLS was changed every 48 h, with vinyl carbamate added to the media every other change, resulting in a cyclic 48 h exposure and 48 h rest cycle. Tissue viability was assessed using a Presto Blue metabolic activity assay every week. At concentrations of 10 and 50 µg/ml vinyl carbamate, hydrogel-embedded PCLS maintained comparable viability to PBS vehicle controls, with relatively high levels of viability (∼80% of starting values) in all groups out to six weeks in culture (Figure 2A). Live/dead staining of hydrogel-embedded PCLS at six weeks in culture revealed that 50-65% of cells were still alive, even within the very center of the construct with no statistical differences between vinyl carbamate and vehicle control exposure (Figure 2B-C). Gene expression of *Cyp2e1*, a critical mediator of vinyl carbamate metabolism, was assessed after six weeks of exposure to vinyl carbamate or PBS. Expression of *Cyp2e1* increased with the higher dose (50 µg/ml) of vinyl carbamate (Figure 2D), which suggested that this dose and dosing scheme had the potential to induce carcinogenesis in hydrogel-embedded PCLS.

**Figure 2:**
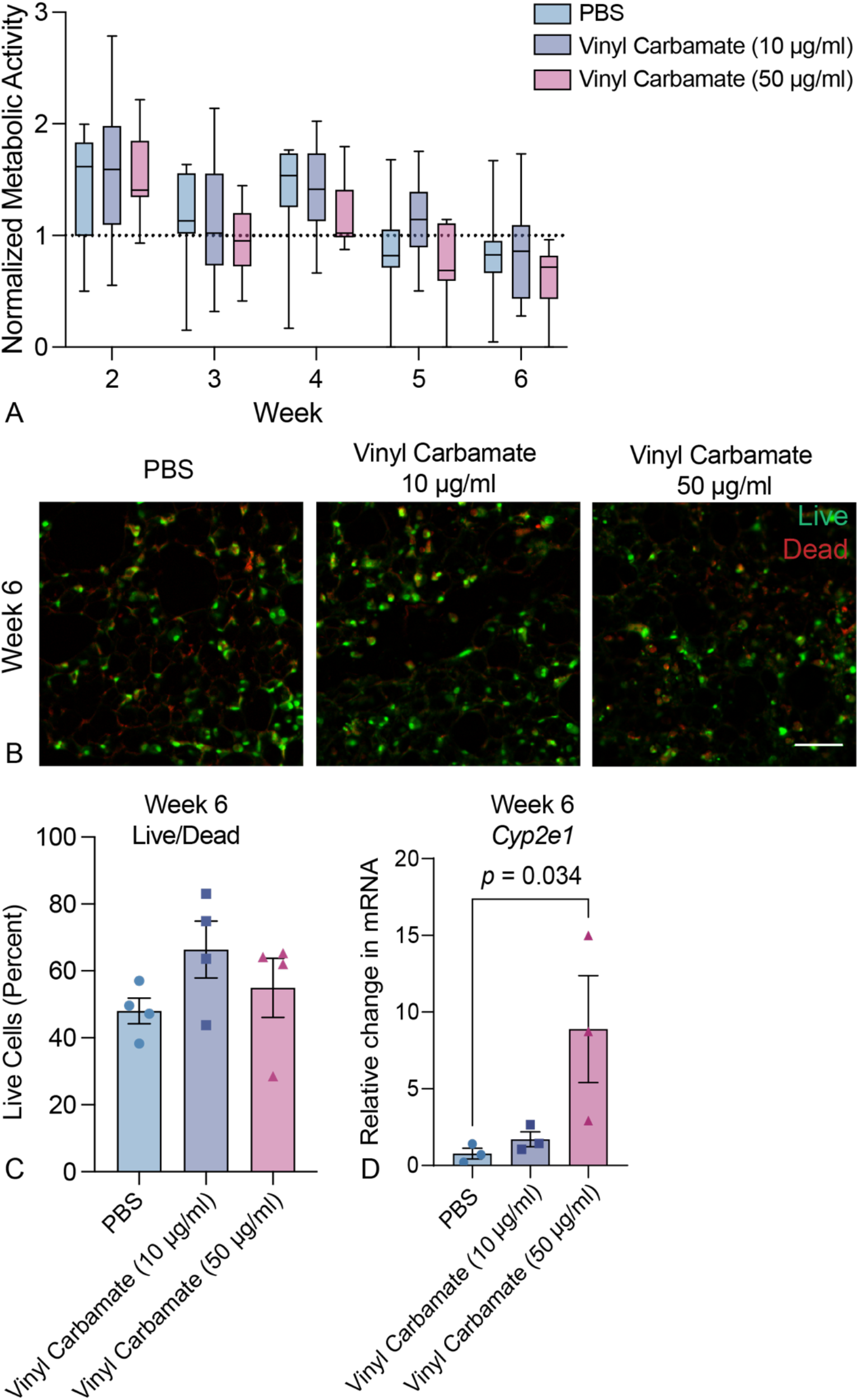
Dose testing showed exposure to 10 or 50 µg/ml vinyl carbamate did not cause overt toxicity in hydrogel-embedded PCLS. A) Presto blue metabolic assessment of hydrogel-embedded PCLS displayed maintenance of viability over six weeks in culture (N=10-11). B) Representative maximum intensity projections from confocal imaging show live (green) and dead (red) staining of hydrogel-embedded PCLS at six weeks. Viability of tissue at the center of the 3D construct was maintained over time. Scale bar, 100 µm C) Quantification of live/dead staining at week six revealed that 50-65% of cells were still alive in all constructs with no statically significant differences (two-way ANOVA; Tukey Test. N=11). D) qPCR analysis showed upregulation of *Cyp2e1* in response to exposure to the higher dose of vinyl carbamate (one-way ANOVA; Kruskal-Wallis test. N=3).

Next, the best hydrogel formulation to support carcinogen-induced premalignancy was determined using the optimal dosing strategy. Four hydrogel conditions were assessed in a design-of-experiments approach: soft with low or high collagen mimic peptide, and stiff with high or low collagen mimic peptide. Proliferation was measured using an EdU incorporation assay, while relevant gene expression changes were quantified using qPCR after six weeks in culture. Proliferation analysis revealed that vinyl carbamate exposure increased the proportion of Edu-positive cells in all hydrogel conditions (Figure 3A, B), with the most robust difference between vinyl carbamate and control conditions observed in stiff hydrogels with a low concentration of collagen mimic peptide. A panel of four genes associated with lung cancer premalignancy were analyzed for expression changes. *Ttf1*, a marker of adenocarcinoma, and *Cox2*, a marker of inflammation and premalignant adenocarcinoma, were anticipated to increase during early adenocarcinoma development. *Apc2* and *Bnip2*, tumor suppressors inactivated in lung cancer, were expected to decrease. PCLS embedded within stiff hydrogels with a low concentration of collagen mimic peptide most adhered to this expected pattern of gene expression (Figure 3C). The proliferation and gene expression data for PCLS embedded within each hydrogel formulation were entered back into the custom statistical design and analyzed with a least-squares regression model to determine the most significant factors and plot the predicted response (JMP Software). The best hydrogel formulation for supporting lung cancer premalignancy in this chemical model was determined to have an elastic modulus of 12.3 kPa and a CGFOGER concentration of 0.1 mM (Supplementary Figure S1).

**Figure 3:**
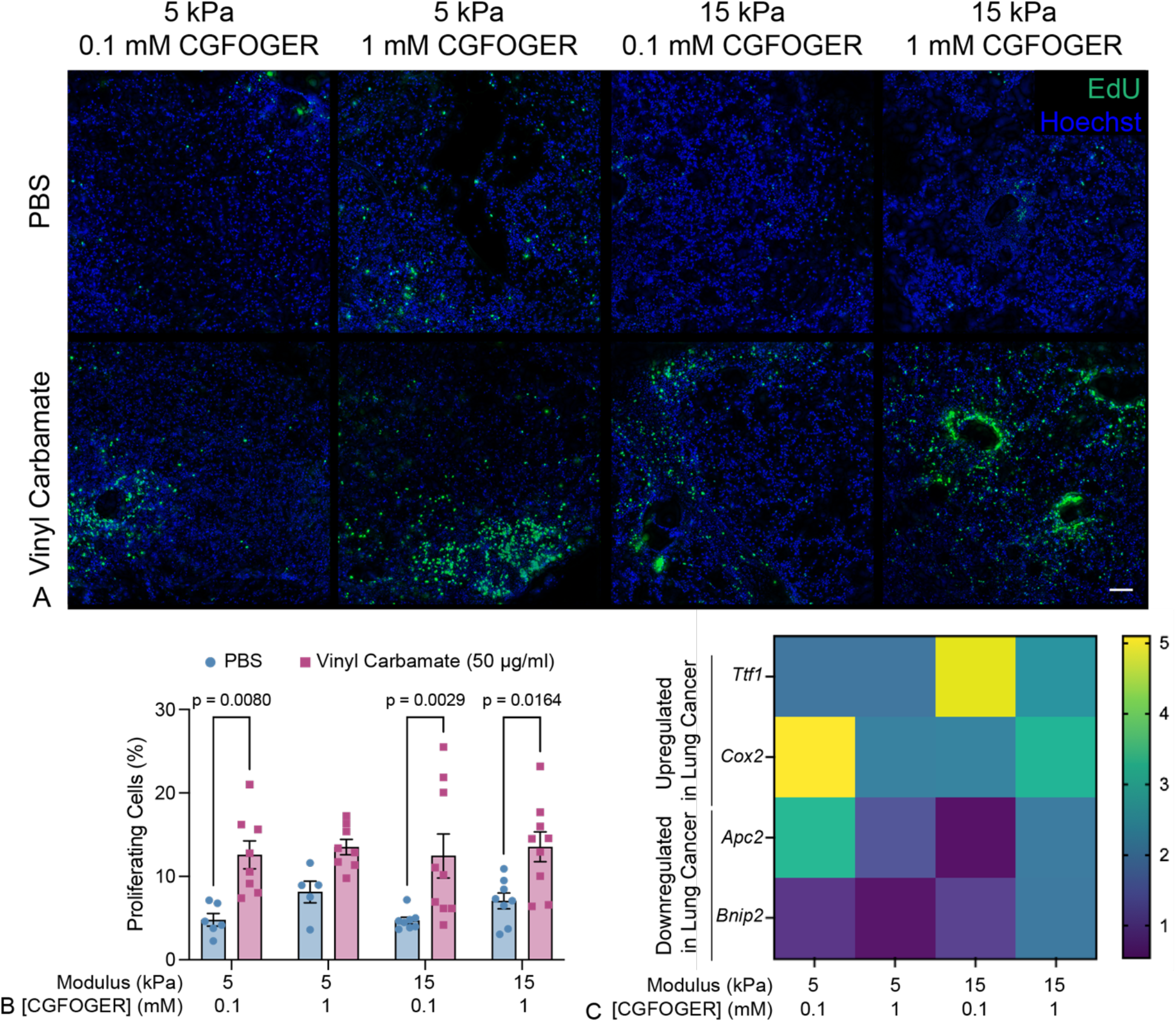
Stiff hydrogels with a low concentration of collagen mimic peptide (GFOGER) provided an environment that supported lung cancer premalignancy in response to vinyl carbamate exposure. A) Representative confocal images (maximum intensity projections) of cellular proliferation within hydrogel-embedded PCLS after six weeks exposure to 50 µg/ml vinyl carbamate. Actively proliferating cells incorporated EdU and were labeled with AlexaFluor488-azide (green). Total nuclei were stained with Hoechst (blue). B) Quantification of the EdU assay revealed increases in proliferation due to vinyl carbamate exposure (N=5-9; two-way ANOVA, Tukey Test). C) Gene expression of analysis of four genes implicated in early lung carcinogenesis showed changes in gene regulation consistent with lung cancer premalignancy in the stiff hydrogel formulation with low GFOGER with vinyl carbamate exposure (N=3).

### Premalignant lesions of mixed phenotype were detected in vinyl carbamate-exposed PCLS

The best hydrogel formulation (stiff with low collagen-mimetic peptide) was selected for PCLS embedding to compare effects of *ex vivo* carcinogen exposure to *in vivo* exposure. Hydrogel-embedded PCLS exposed to 50 µg/ml vinyl carbamate for six weeks were assessed for tissue morphology, tissue stiffness, gene expression, and cellular content. Vinyl carbamate and PBS-exposed hydrogel-embedded PCLS sections were H&E stained and assessed by an expert lung pathologist. The pathologist observed regions of macrophage infiltration, alveolar atypia, sub-pleural fibrosis, and bronchial dysplasia (Figure 4A) only in the vinyl carbamate-exposed samples. Since increases in tissue stiffness are concurrent with tumor development *in vivo*, atomic force microscopy (AFM) was used to assess the relative elastic modulus of discrete regions of abnormal tissue morphology in vinyl carbamate-exposed PCLS. AFM was also performed on healthy control lungs and premalignant lesions generated *in vivo* by exposure to urethane, the metabolic precursor of vinyl carbamate. Premalignant regions generated by *in vivo* chemical carcinogen exposure were notably stiffer than healthy lung tissue. Interestingly, *ex vivo* lesions tended to be even stiffer (Figure 4B), and exhibited a greater variability in degree of stiffness, even in regions of similar topography (Figure 4C, D). This enhanced stiffness may be due to an abundant accumulation of fibrillar collagen, as assessed by picrosirius red staining (Supplementary Figure S2). Expression of a panel of genes shown to change in response to urethane exposure *in vivo*^11,54^ and expected to change in response to vinyl carbamate treatment revealed changes consistent with premalignancy, including significant upregulation of *Il6*, *Il1β*, *Ttf1*, and *Ubac2*, as well as downregulation of *Apc2*, *Ecad*, and *Ces1*. *Cox2* had a non-significant increase with vinyl carbamate compared to control. Only one of the genes assessed, *Bnip*, did not show significantly different expression in response to vinyl carbamate exposure (Figure 4E). Overall, these data suggest that *ex vivo* exposure of PCLS to vinyl carbamate resulted in a variety of pathological traits consistent with premalignant lesion development.

**Figure 4:**
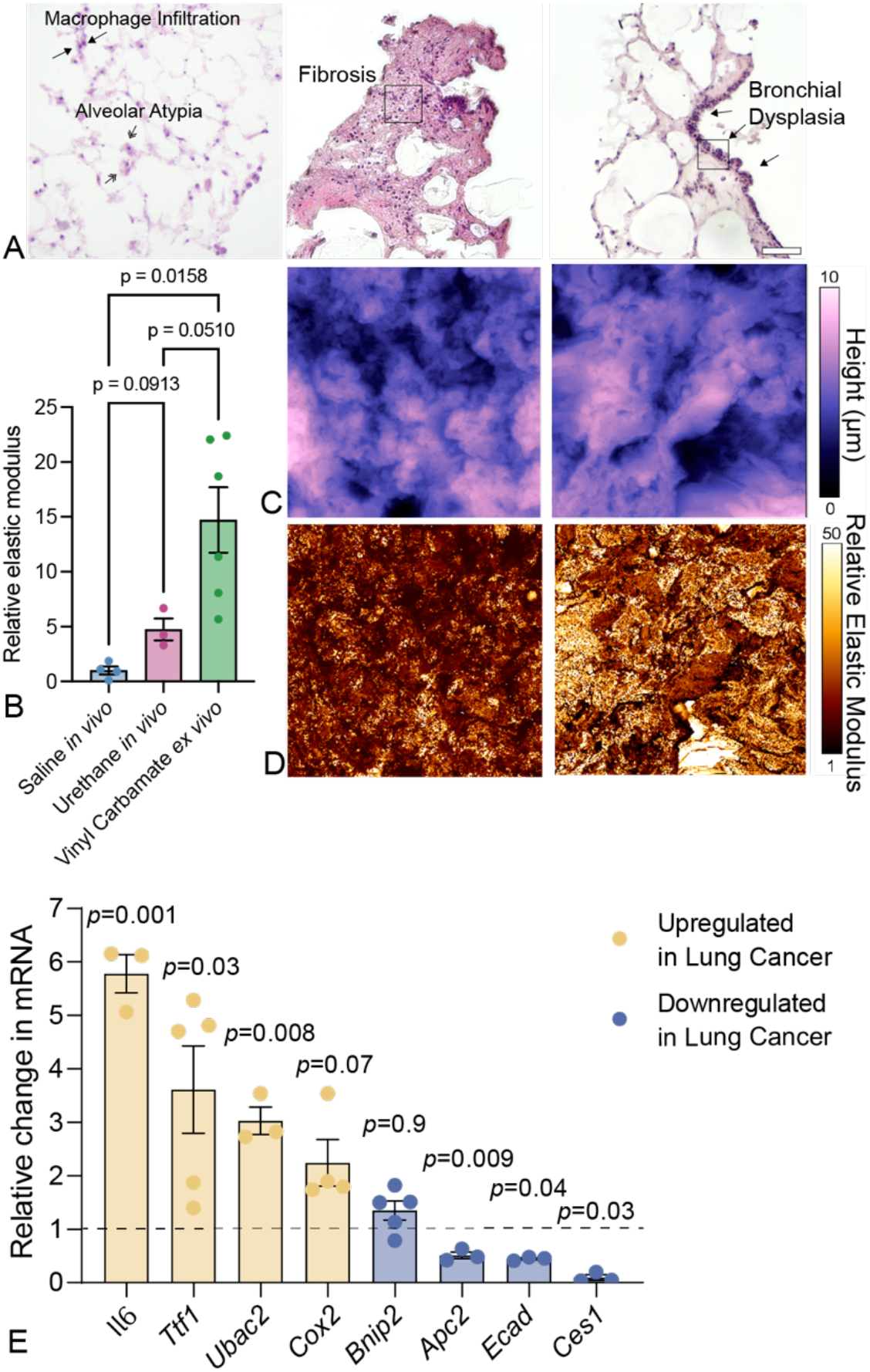
Lesions formed in the tissue-engineered models recapitulated characteristic markers of premalignancy. A) Representative H&E images of hydrogel-embedded PCLS following six weeks of exposure to 50 µg/ml vinyl carbamate in the best hydrogel formulation. Regions of macrophage infiltration (left, single-headed arrows), alveolar atypia (left, double-headed arrows), fibrosis (center), and bronchial dysplasia (right, single-headed arrows) were identified by a pathologist. Boxes indicate regions assessed by AFM. Scale bar, 100 µm. B) AFM analysis showed that both *in vivo* and *ex vivo* premalignant lesions had high stiffness relative to normal lung tissue. C) Topography maps of the 50 x 50 µm regions identified by boxes in panel A showed variable tissue heights up to 10 µm. D) Stiffness maps of the same regions showed the variability in relative elastic modulus between *ex vivo* premalignant lesions. E) qPCR analysis of six genes implicated in lung carcinogenesis revealed gene expression patterns consistent with premalignancy observed *in vivo* (two-tailed T-test; N=3-5).

Additional immunofluorescence staining was performed to more closely examine the cellular content and pathologies induced by vinyl carbamate exposure in PCLS after six weeks in culture. In all conditions, staining for E-cadherin and vimentin revealed the persistence of both epithelial and mesenchymal cells over this extended time in culture (Supplementary Figure S3A). Alveolar type 1 and type 2 epithelial cells were detected (Supplementary Figure S3B). Neutrophils were not detected (Supplementary Figure S3C). Staining for CD68 showed the persistence of macrophages in PCLS and suggests an accumulation of macrophages in response to vinyl carbamate exposure (Figure 5A). This is congruent with the observation of macrophage infiltration by H&E staining (Figure 4A). A small population of T-cells (CD3^+^) was also present after six weeks (Figure 5A). PCLS tissue lacks a vascular supply of blood-derived monocytes. Therefore, any increase in macrophages over time should be due to local proliferation rather than recruitment. Co-staining of CD68 and Ki67 confirmed many proliferative macrophages in vinyl carbamate-exposed PCLS (Figure 5B). Additional immunofluorescence staining provided further characterization of the bronchial and alveolar abnormalities noted on H&E images. Staining for CYP2E1 highlights potential metabolism of vinyl carbamate. Keratin 5 (KRT5) marks an early squamous cell phenotype and TTF1 marks an early adenocarcinoma phenotype. CYP2E1+ cells were sparse and scattered in the tissues exposed to the vehicle control. TTF1 was not detected in the PBS vehicle control, but there was a low level of KRT5 expression present along the airways of this tissue (Figure 5C). Given that KRT5 is not typically found in lower airways of healthy murine lung^55^, this observation may suggest a cellular response to injury during slicing or maintenance in long-term culture^56^. In vinyl carbamate exposed PCLS, particularly around the regions of prominent bronchial dysplasia, there was an increase in CYP2E1+ cells, as well as notable expression of both KRT5 and TTF1 (Figure 5C), suggesting that *ex vivo* lesions display a mixed pathological phenotype rather than the predominant adenocarcinoma phenotype produced by vinyl carbamate *in vivo*.

**Figure 5:**
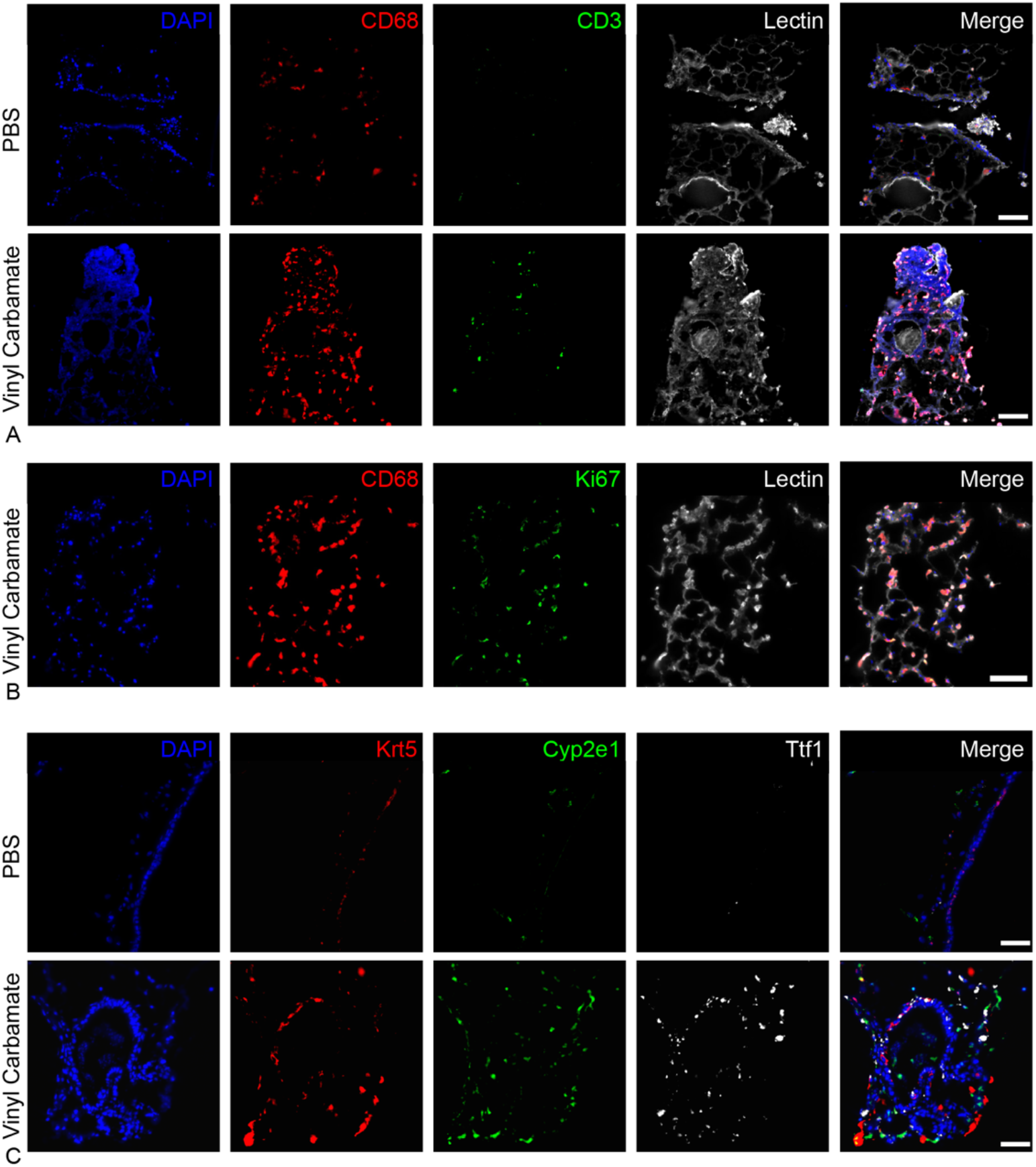
Exposure to vinyl carbamate alters immune and cancer cell markers in hydrogel-embedded PCLS. A) Representative immunofluorescent images stained for CD3 (T-cells), CD68 (macrophages), and Lectin (tissue structure) showed an accumulation of macrophages in vinyl carbamate-exposed, hydrogel-embedded PCLS. B) Representative immunofluorescent images stained for Ki67 (proliferation), CD68 (macrophages), and Lectin (tissue structure) demonstrated active proliferation of macrophages. C) Representative immunofluorescent images stained for CYP2E1 (vinyl carbamate metabolism), KRT5 (squamous carcinoma), and TTF1 (adenocarcinoma) revealed regions of mixed premalignant phenotype in vinyl carbamate-exposed, hydrogel-embedded PCLS. Scale bars, 100 µm.

### Chemoprevention agents acted on hydrogel-embedded PCLS

Both an established and an emerging chemoprevention agent penetrated the hydrogel embedding layers and induced gene expression changes protective against tumorigenesis in hydrogel-embedded PCLS. Iloprost is a lung cancer chemoprevention agent that produces an anti-cancer effect in part by increasing PPARγ expression and activity and has demonstrated efficacy on bronchial dysplasia in a clinical trial.^11,42,57-59^ Curcumin is extracted from the turmeric plant and has anti-inflammatory and growth suppressive effects in lung cells and mice.^60-62^ First, hydrogel-embedded PCLS were treated with 10 μM iloprost in culture and metabolic activity was assessed by Presto Blue analysis on days 0, 3, and 7, to confirm that iloprost did not affect viability of hydrogel-embedded PCLS. Results showed no significant changes in metabolic activity over 7 days (Figure 6A). Iloprost induced a trend towards increased *Ppar*γ mRNA expression by qPCR (Figure 6B), and a significant increase in PPARγ activity as assessed by a dual-luciferase PPRE assay (Figure 6C) in hydrogel-embedded PCLS. Viability analysis revealed that 3.0 µg/mL curcumin did not exhibit cytotoxicity in hydrogel-embedded PCLS (Figure 6D). qPCR analysis showed that curcumin induced significant upregulation of *Ppar*γ expression in conjunction with decreased *Il6* expression (inflammation) (Figure 6E). Together these data suggest that the hydrogel embedding does not act as a significant barrier to drug delivery verifying that tissue-engineered models of lung cancer premalignancy will be powerful tools for chemoprevention screening and validation.

**Figure 6:**
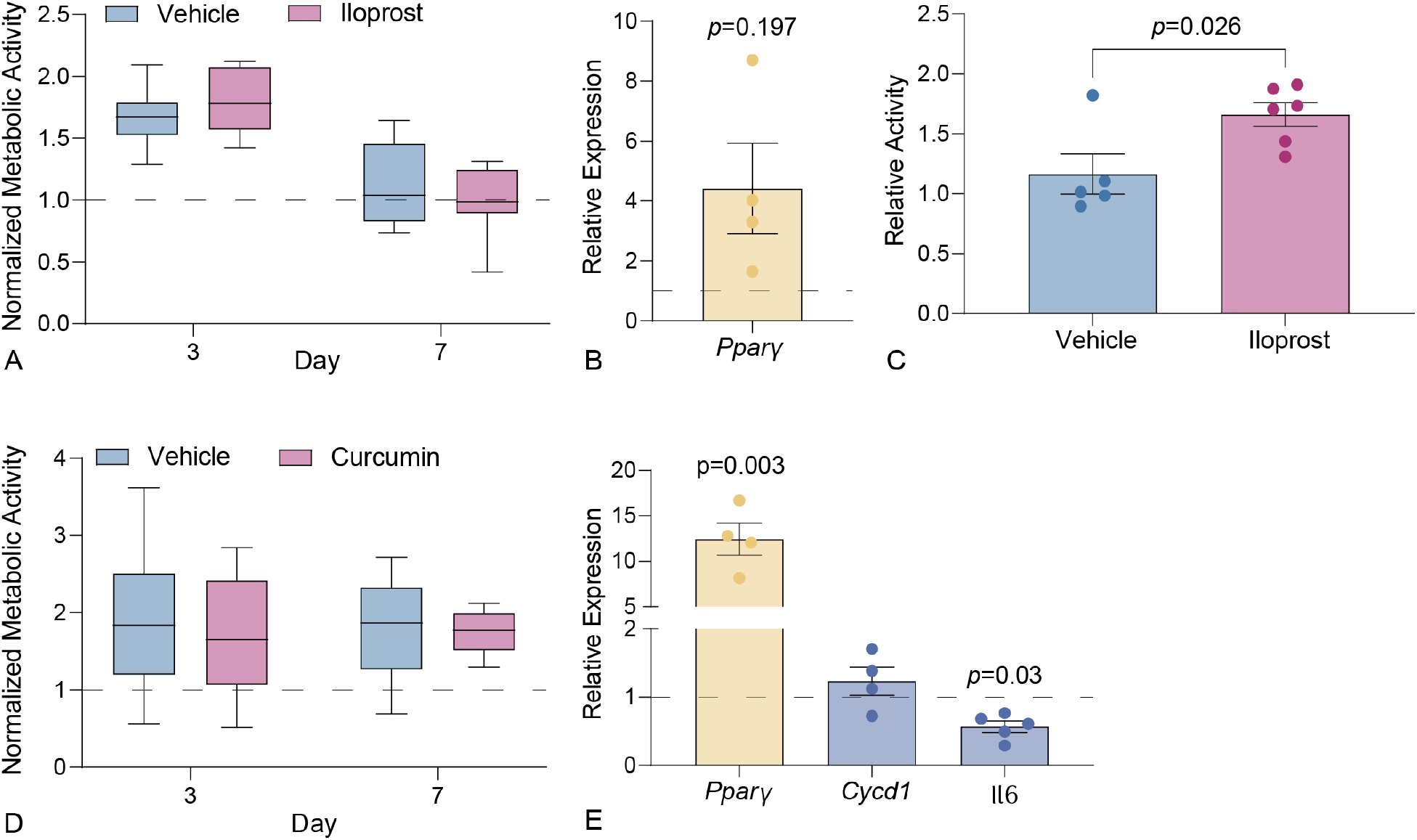
Chemoprevention agents diffused through hydrogels and act on hydrogel-embedded PCLS. A) A Presto Blue assay for metabolic activity monitored viability of hydrogel-embedded PCLS at day 0, 3, and 7 during treatment with a vehicle control or 10 μM iloprost. Data are shown relative to day 0 (dotted line). There were no statistical differences between vehicle and treatment (two-way ANOVA; Tukey Test; N = 8-10). B) Expression of *Ppar*γ was measured by qPCR at day 7 and presented relative to vehicle control (dotted line; two-tailed T-test, N=4)). C) Activity of PPARγ was increased on day 7 (two-tailed T-test; N=5). D) A Presto Blue assay measured viability of PCLS at day 0, 3, and 7 during treatment with vehicle or 3 μg/ml curcumin. Data are shown relative to day 0 (dotted line). There were no statistical differences between vehicle and treatment (two-way ANOVA; Tukey Test; N=12.) E) Expression levels of curcumin responsive genes were measured by qPCR at day 7 and shown relative to vehicle control (dotted line; two-tailed T-test, N=4-5).

### Tissue-engineered lung model parameters were validated in human hydrogel-embedded PCLS

Human PCLS were incorporated into our tissue-engineered model of lung cancer premalignancy to validate that the conditions selected using murine tissues could also produce premalignant phenotypes in a human model. Human PCLS were either embedded in the stiff hydrogel with low collagen-mimetic peptide concentration or cultured without hydrogel (non-embedded) for six weeks. Non-embedded human PCLS showed decreasing metabolic activity over time with a trend toward larger decreases overall than hydrogel-embedded PCLS over the course of six weeks. Vinyl carbamate exposure did not cause a significant reduction in viability (Figure 7A). Proliferation was assessed by EdU assay and gene expression profiled in hydrogel-embedded human PCLS after six weeks in culture. As with murine tissue, exposure of human PCLS to vinyl carbamate resulted in an overall increase in proliferation (Figure 7B). The expression of multiple genes also showed changes consistent with the development of premalignant lesions *in vivo. UBAC2*, *COX2*, and *CYP2E1* expression increased while *BNIP2* and *CES1* decreased (Figure 7C). In contrast to the murine results, *TTF1* expression levels did not change, perhaps suggesting an even higher predisposition to squamous malignancy in human PCLS.

**Figure 7:**
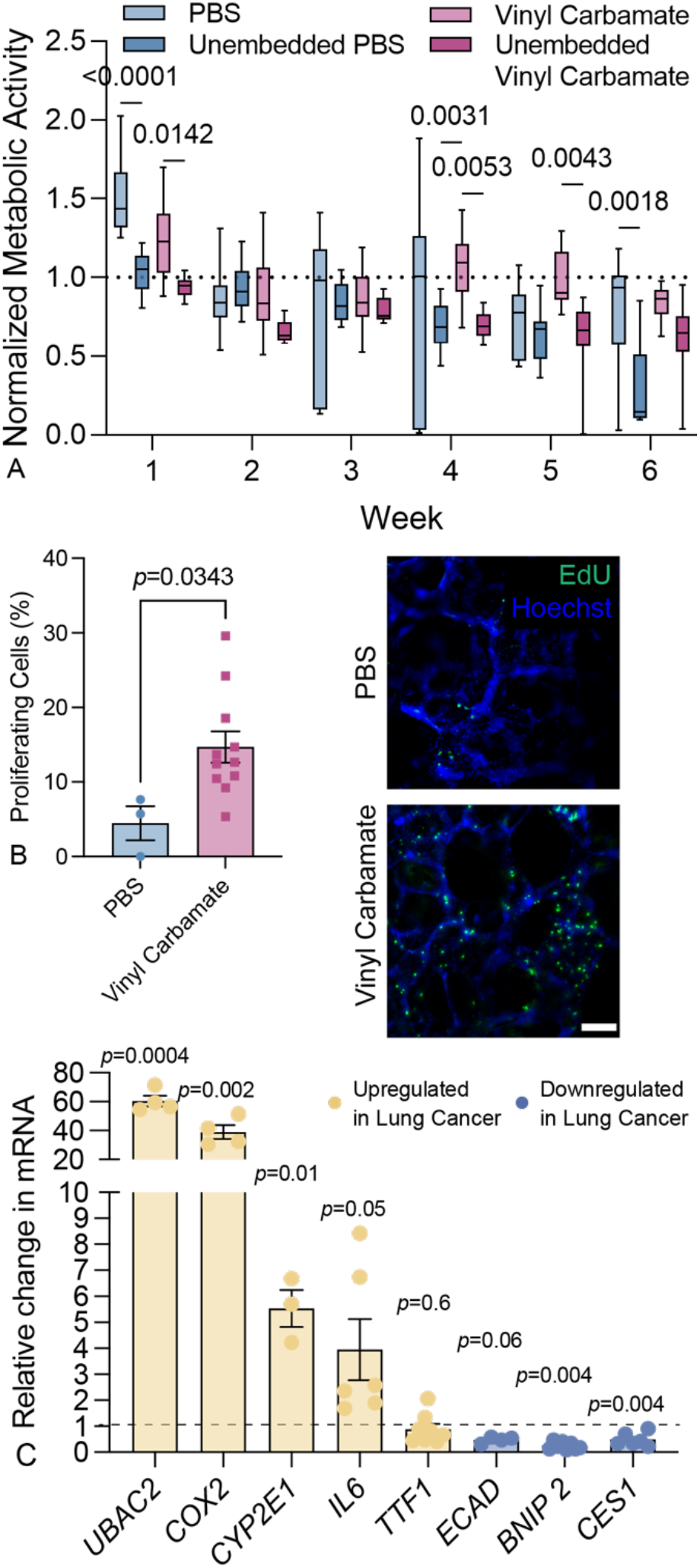
Human, hydrogel-embedded PCLS exposed to vinyl carbamate developed premalignant phenotypic changes over six weeks in culture. A) A Presto Blue assay measured viability of hydrogel-embedded or unembedded PCLS weekly through day 42 during exposure to vehicle or vinyl carbamate (50 μg/mL). Unembedded PCLS showed reduced viability relative to embedded counterparts at multiple timepoints (two-way ANOVA, Tukey test; N=6-18. Data are shown relative to day 0 (dotted line) B) EdU assay demonstrated increased proliferation in human PCLS as a response to vinyl carbamate exposure (two-tailed T-test, N=3-12; Scale bar, 200 µm). C) Expression of genes expected to be altered by vinyl carbamate was measured by qPCR at day 42, showing consistency with premalignant phenotype (two-tailed T-test; N=3-5).

## Discussion

This study describes a strategy for engineering biomimetic model of early lung carcinogenesis using hydrogel-embedded murine PCLS and validation of the design parameters with human tissue. PCLS contain a multitude of cell types and extracellular components preserved in the native pulmonary microarchitecture, which is a critical advantage when studying complex pathologies such as lung cancer. Many pioneering *in vitro* studies of lung cancer have employed sophisticated organoid cultures^63-66^. While this approach provides a 3D microenvironment, it typically consists of only one or two cells types or lineages, and is limited in the complexity of the geometry or maturity of the ECM that cells produce^67^. PCLS, in contrast, display high levels of complexity, and until recently^39^ have been primarily limited by short-term viability and functionality *ex vivo*^12^. This study employs an engineering approach to embed murine and human PCLS within rationally designed hydrogel biomaterials to preserve the longevity and utility of PCLS.

The hydrogel formulations evaluated in this study were designed to replicate the contributions of collagen accumulation and thus increased microenvironmental stiffness during early-stage carcinogenesis. Studies of lung cancer patient tissues have shown that *in vivo*, dense ECM is a contributor to tissue stiffness, and these linked phenomena enhance tumor cell proliferation and invasive phenotype^68^. Dense fibrillar collagen promotes proliferation of epithelial cells, healthy or cancerous, by activation of integrin-mediated FAK signaling^34,69^. Fibrillar collagen has also been implicated in tumor invasiveness during metastatic transition^33,70^, making collagen dynamics a critical aspect for modeling premalignancy. One limitation of the use of a CGFOGER peptide in our studies is that it lacks the full architecture of fibrillar collagen, which has been proven to both change with and directly promote tumor progression^33,70^. Future studies with hydrogel-embedded PCLS could use hybrid-hydrogels that incorporate decellularized ECM (dECM) containing larger intact proteins and growth factors^71-73^. A dynamically stiffening hybrid-hydrogel would also allow decoupling of collagen accumulation and stiffness to assess relative individual contributions of these phenomena in a way not possible *in vivo.* Other aspects of the hydrogel formulation could also be modified to recapitulate specific aspects of tumor biology. The synthetic crosslinker employed here could be fully or partially replaced with an MMP-degradable sequence, allowing for cells to locally remodel the hydrogel^74^. MMP expression generally increases with cancer progression, and different cancer subtypes can be characterized by the expression of specific MMPs^75,76^. Targeted MMP-degradable sequences could, therefore, provide an environment specifically tuned to support the development of a specific cancer subtype. The adhesion peptides used could be further expanded or optimized, based on data about which ECM components are critical during tumorigenesis. For example, non-fibrillar collagens and other structural ECM components such as Tenasin C also play important roles in lung cancer pathogenesis, again with differential ECM signatures present in different cancer subtypes^77,78^. However, systematic profiling is needed to determine how ECM composition changes in premalignancy and during progression of lung lesions into tumors. Alternatively, instead of targeted crosslinker sequences or adhesion peptides, a dECM hybrid-hydrogel incorporating material from either healthy or diseased lung could mimic the full spectrum of ECM changes that occur during cancer progression^72,73^. The hydrogel-embedded PCLS model can be the foundation for a host of future studies exploring interactions between lung lesions and the surrounding matrix.

Here, we demonstrated induction of premalignant phenotypes with *ex vivo* exposure of tissues to a tobacco carcinogen. Hydrogel-embedded PCLS displayed a mixture of various histologies in response to vinyl carbamate exposure, including limited regions of alveolar atypia, enhanced macrophage proliferation, fibrosis, and squamous bronchial dysplasia. In mice, vinyl carbamate and its metabolic precursor, urethane, induce adenocarcinomas, following a predictable pattern of progression of early hyperplasia to carcinoma^79,80^. The mixed histology produced in PCLS by vinyl carbamate was therefore unexpected. One potential explanation of this observation relates to bioavailability. In PCLS, internal regions of the lung are exposed during slicing, and the exposure protocol covered the entire PCLS in media containing carcinogen. This procedure could result in the carcinogen targeting cells that might be less exposed *in vivo*. CYP2E1, the primary enzyme which metabolizes vinyl carbamate, is expressed at the highest levels in the lung in club cells, with lower expression in alveolar epithelial type 2 cells^81^. If vinyl carbamate reaches club cells in the airway, metabolizing mechanisms are in place that could lead to unexpected carcinogenesis in PCLS.

Likewise, in our study, low-level expression of KRT5, a basal cell marker, was observed in airway epithelium of vehicle control (PBS)-treated hydrogel-embedded PCLS after six weeks in culture. In contrast to humans, KRT5 staining and basal cells are normally present only in mouse trachea, not lung.^55^ Long-term culture of hydrogel-embedded PCLS may produce basal cell phenotypes, and subsequent carcinogen exposure could induce progression towards squamous bronchial dysplasia. A prior study of long-term culture of non-embedded PCLS observed non-malignant keratinizing squamous metaplasia develop over the course of 28 days in culture^56^. In contrast to our study, the PCLS in this study were generated from either fibrotic or tumor-bearing human lungs, conditions which may predispose even non-involved tissue to altered pathology. In addition, all the PCLS had areas of altered histology, though it was not considered cancer-related, while in hydrogel-embedded PCLS we did not observe histological changes pronounced enough to be observed in H&E-stained samples. These results nevertheless suggest that long-term culture itself could change tissue morphology; however, hydrogel-embedding appears protective to some degree. Another important consideration for any PCLS study is that slicing is a source of tissue damage which will undoubtedly cause localized healing responses and potentially contribute to the appearance of fibrotic areas over extended periods in culture as observed in the murine studies reported here. Lung fibrosis and lung cancer share a number of cellular level traits, including enhanced fibroblast activation and ECM accumulation^82^. Idiopathic pulmonary fibrosis (IPF) is also a strong risk factor for the development of lung cancer, with a tendency to predispose towards squamous cancer^83^, and the ECM signature of squamous cancer shows a great deal of overlap with that of fibrosis^78^. In regions identified in this study as bronchial dysplasia by morphology, cells expressing both KRT5—a squamous carcinoma marker—and TTF1—an adenocarcinoma marker—were observed, suggesting that application of vinyl carbamate *ex vivo* yields mixed-histology premalignant lesions. Treatment of hydrogel-embedded PCLS with other carcinogens may produce more robust models of specific histology. Future studies could explore a range of carcinogens and exposure protocols to optimize generation of premalignant lesions that recapitulate *in vivo* models.

Prior studies of chemical carcinogenesis using 3D cell culture models have largely focused on epithelial-cell intrinsic pathologies. Rayner et. al. cultured primary normal human epithelial cells at an air-liquid interface (ALI) using transwell inserts and exposed the resulting airway-mimic epithelial layers to cigarette smoke extract (CSE). The results showed increased proliferation resulting in a thickening of the epithelial cell layers^84^, potentially analogous to the bronchial dysplasia observed in our study. Using a similar methodology, Eenjes et. al. noted upregulation of oxidative stress pathways in primary epithelial cell layers treated with CSE, along with increased secretion of IL-8. While they noted that a pro-inflammatory phenotype was expected in response to CSE exposure, the model did not contain any inflammatory cells for studying the downstream effects of IL-8^85^. Co-culture models can provide valuable insights into cellular crosstalk during disease. For example, a study that bioprinted a mixture of cancer cell and fibroblast lines into a 3D hydrogel exposed cells in this culture system to CSE. This model resulted in the acquisition of EMT markers by the cancer cell line, while treatment with an established chemotherapeutic showed a decrease in viability^86^. Organoids have also proven to be a very powerful tool for multicellular models of cancer. Primary tumor-derived organoids containing lung cancer cells alongside both fibroblasts and inflammatory cells showed a higher resistance to chemotherapeutics than did primary cells grown on tissue-culture plastic, highlighting again the importance of environmental context in predicting cellular responses to treatment^87^. While each of these reductionist models has their uses, they all recapitulate only a limited range of cellular interactions, e.g., all airway cells with no model of the lung parenchyma, pre-selected and mixed cell lines, or all cells derived from established, late-stage tumors. In this study, we present a highly multicellular model of the earliest stages of cancer development which could be of use in studying both the complex biology of tumorigenesis and integrated cellular responses to cancer treatments.

A major goal of studying premalignant lung lesion biology is to identify and validate interventions that reduce the risk of lung tumors. *In vitro* or organoid models can identify effects of chemoprevention on specific cell types or interactions. However, models that recapitulate complex responses observed *in vivo* are required to understand the mechanisms of chemoprevention agent interception of premalignant lesions. Chemoprevention studies rely on *in vivo* models for evidence of efficacy before moving to clinical trials. Hydrogel-embedded PCLS models of lung cancer premalignancy offer an intermediate step between *in vitro* and *in vivo* chemoprevention studies. We demonstrated this by treating hydrogel-embedded PCLS with two chemoprevention agents and validating similar responses compared to *in vivo* models.

These studies were conducted for only seven days, but since metabolic activity and signaling continues beyond seven days, we expect that the responses would continue throughout the six-week model. While agents that rely on recruited immune cells or enzymes that are not present in PCLS may not be effective, many agents are expected to be active in PCLS and this model could screen agents for efficacy before moving to full *in vivo* testing. The opposite may also be true, that some agents may work in PCLS but be ineffective in the context of a full immune system, however overall, we expect that *ex vivo* testing of agents in PCLS would reduce the number of animals used for chemoprevention screening and mechanism studies. Decreasing cost and stress will improve feasibility of chemoprevention studies, increasing the opportunities for translation of agents to clinical application.

Models of lung cancer premalignancy were engineered by embedding thin slices of lung tissue within rationally designed hydrogel biomaterials. Both carcinogens and chemoprevention agents were easily supplemented into the culture media and diffused through the hydrogel to influence the embedded tissue. Exposure of hydrogel-embedded PCLS to the tobacco smoke carcinogen vinyl carbamate induced a variety of pathologies including alveolar atypia, macrophage infiltration, fibrosis, and bronchial dysplasia. The design parameters established using murine tissue were validated with human PCLS demonstrating that tissue-engineered models of lung cancer premalignancy are a promising new tool that will improve our understanding of cancer biomechanics, ECM dynamics, carcinogenesis, lung lesion progression, and chemoprevention.

## Acknowledgements

The authors thank Dr. Kolene Bailey for contributing Figure 1E as well as Dr. Hong Wei Chu and Niccolette Schaefer for providing agarose-filled, de-identified human lung samples. This study was supported by funding from the National Cancer Institute (R21 CA252172 to RB, MT, and CMM), NIH (K08HL155894 to PSH, K25HL148386 to BP, and 5T32HL007085-47 to RB), the Ludeman Center for Women’s Health (seed grant to BP) and by generous grants of the John Patrick Albright (to BP).

## Ethics Declaration

The authors declare no competing interests.

## Source Data

The raw and processed data required to reproduce these findings are available here: Blomberg, Rachel; Sompel, Kayla; Hauer, Caroline; Pena, Brisa; Driscoll, Jennifer; Hume, Patrick; Merrick, Daniel; Tennis, Meredith; Magin, Chelsea (2023), “Tissue-engineered models of lung cancer premalignancy”, Mendeley Data, V1, doi: 10.17632/gm93j5r9fw.1.

