## Supplementary Information for "Tissue-engineered models of lung cancer premalignancy"

| Primer | Unique Assay ID |
| --- | --- |
| TTF1 | Hu: qHsaCID0009451<br>Mu: qMmuCED0046157 |
| COX2 | Hu: qHsaCID0020933.<br>Mu: qMmuCED0003742. |
| APC2 | Hu: qHsaCID0013039<br>Mu: qMmuCED0004889 |
| IL6 | Hu: qHsaCID0020314<br>Mu: qMmuCED0045760. |
| UBAC2 | Hu: qHsaCID0007335<br>Mu: qMmuCID0017311 |
| CES1 | Hu: qHsaCID0021826.<br>Mu: qMmuCID0026391 |
| IL1b | Hu: qHsaCED0044677<br>Mu: qMmuCID0005641 |
| E-CAD | Hu: qHsaCID0015365<br>Mu: qMmuCED0044197 |
| CYP2e1 | Hu: qHsaCID0018067<br>Mu: qMmuCID0019056 |
| PPARγ | Hu: qHsaCED0044425<br>Mu: qMmuCID0018821 |
| BNIP2 | Hu: qHsaCID0009133<br>Mu: qMmuCED0044170 |
| VIM | Hu: qHsaCED0042034<br>Mu: qMmuCED0046651 |
| TNFα | Mu: qMmuCED0004141 |
| NFkB | Mu: qMmuCID0005357 |
| CycD1 | Mu: qMmuCID0023518 |

Table S1: Biorad Prime PCR Assays. The unique ID of each primer set for human (hu) and murine (mu) assays.

| Target | Antibody | Host | Dilution |
| --- | --- | --- | --- |
| CD3 | Biolegend, 100206 | Rat | 1:50 |
| CD68 | Biolegend, 137002 | Rat | 1:500 |
| Cyp2e1 | Thermo Fisher, MA525001 | Mouse | 1:100 |
| E-cadherin | ProteinTech, 20874-1-AP | Rabbit | 1:100 |
| Keratin 5 | BioLegend, 905901 | Chicken | 1:200 |
| Ki67 | Thermo Fisher, | Rat | 1:250 |
| Ly6G | BD Biosciences | Rat | 1:50 |
| PDPN | Thermo Fisher, MA5-18054 | Hamster | 1:100 |
| SFTPC | Thermo Fisher, PA571680 | Rabbit | 1:50 |
| Vimentin | Thermo Fisher, MA511883 | Mouse | 1:00 |

Table S2: Primary Antibodies. The target, source, host species, and concentration of each primary antibody used for IF analysis.

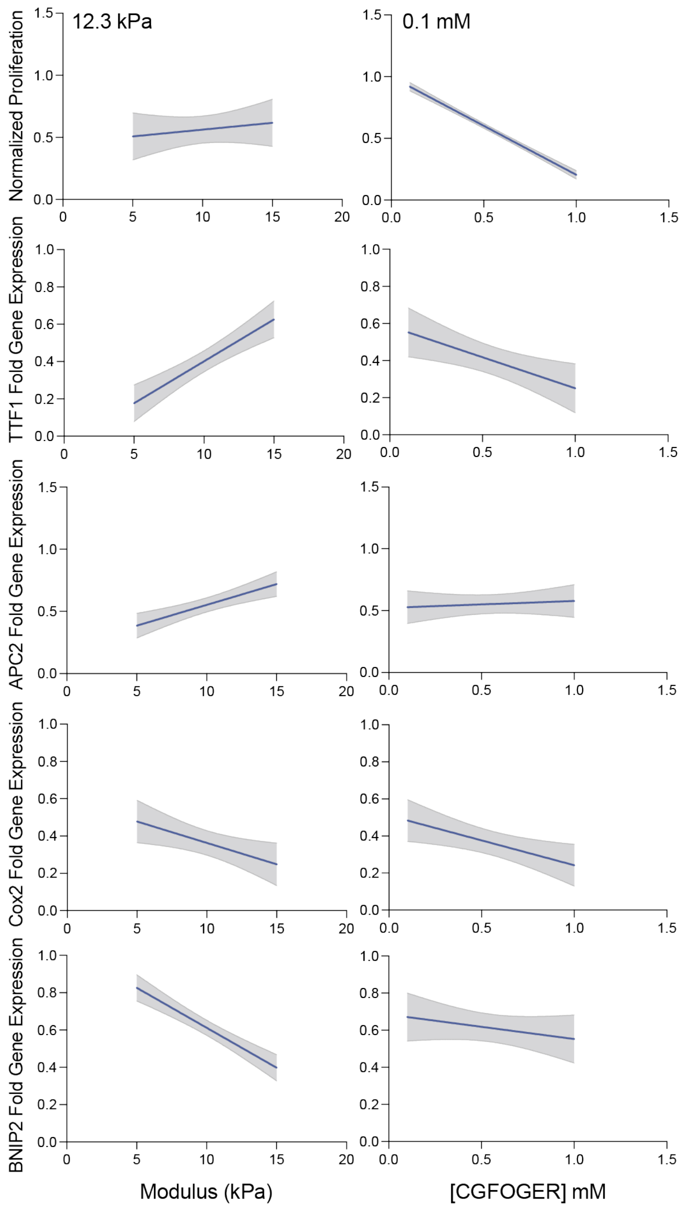


Figure S1: Design-of-experiment analysis for optimal hydrogel formulation. The results of EdU staining and qPCR from four hydrogel formulations were input to JMP software with the most desirable results being a strong difference between vehicle and vinyl carbamate treatment in proliferation and gene expression.

| Slide | Comment |
|---|---|
| StLo VC 3-19 | Areas of septal fibrosis most prominent in the subpleural area; Macrophages present, intra-alveolar; Area of bronchial dysplasia. Maybe a few atypical pneumocytes but no adenoma or even early AAH like change |
| StLo VC 3-7 | Maybe a little bit of bronchial dysplasia |
| StLo VC 2-19 | Some atypical alveolar lining cells in sections near the slide label – don't think these are macrophages; No airways in these sections |
| StLo VC 2-7 | Some macrophages, maybe rare mildly atypical pneumocyte; No major alveolar septal changes; No airways present |
| StLo VC 1-19 | Septal fibrosis, mild to moderate parenchymal foci (2); Mixed inflammatory cells both interstitial and intra-alveolar with macrophages (mostly intra alveolar), neutrophils and lymphocytes/plasma cells (mostly interstitial); significant pneumocyte atypia; No airways |
| StLo VC 1-7 | Scattered macrophages; Airway without abnormality present; No septal fibrosis |

Table S4: The complete pathology report detailed various abnormalities in vinyl carbamate (VC) treated PCLS only.

**Picrosirius Red Staining Methods**

Frozen sections were allowed to thaw at room temperature and then dipped in distilled water to clear OCT. Slides were submerged in picrosirius red solution (Abcam) for 60 min, washed twice in 0.5% acetic acid, and then dehydrated by subsequent submersion in 100% Ethanol (2 x 5 min) and SafeClear II (2 x 5 min, Fisher Scientific). Slides were mounted in Cytoseal 60 (ThermoFisher) under a 24 x 50 mm coverslip and then imaged on a brightfield microscope (Olympus BX63)

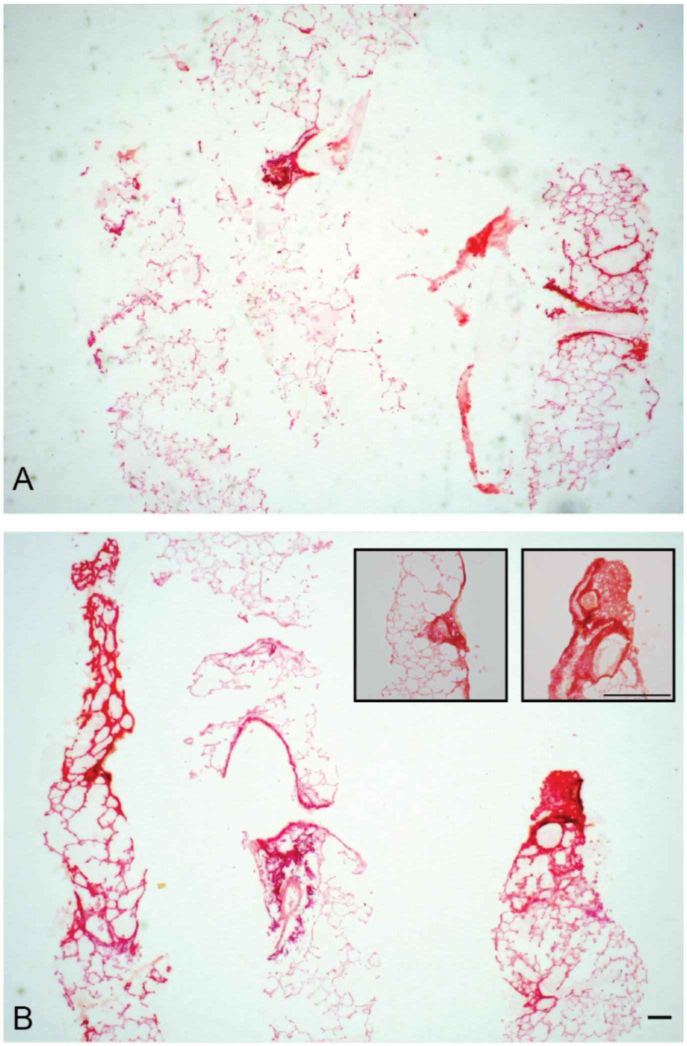


Figure S2: Picrosirius Red Stain of both A) PBS and B) vinyl carbamate treated PCLS showed accumulation of fibrillar collagen after vinyl carbamate treatment. Insets show regions of interest identified in the pathology report. Scale bars, 150 μm.

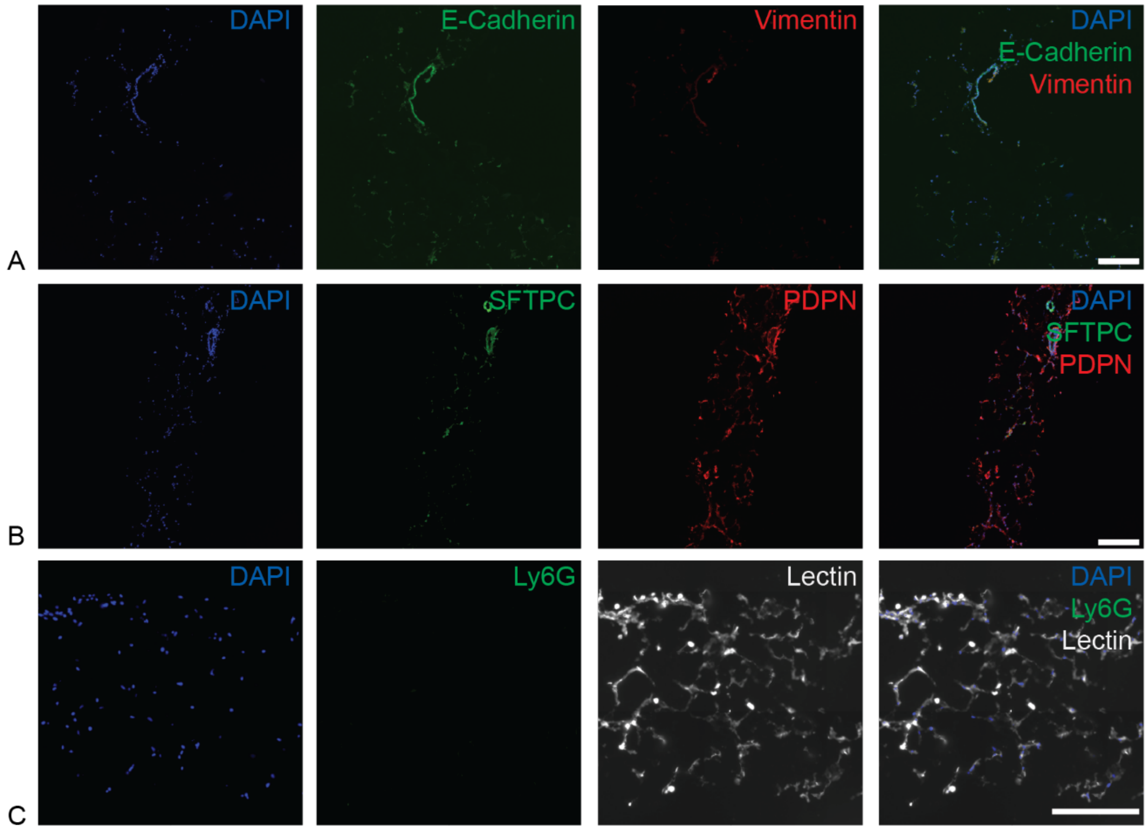


Figure S3: Immunofluorescence staining after six weeks in culture showed the persistence of multiple relevant lung cell populations, including A) E-cadherin$^{+}$ epithelial cells and vimentin$^{+}$ fibroblasts, and B) PDPN$^{+}$ Alveolar Type I cells and SPC$^{+}$ Alveolar Type II cells. C) Ly6G$^{+}$ neutrophils were not detected.
